# scmTE: multivariate transfer entropy builds interpretable compact gene regulatory networks by reducing false predictions

**DOI:** 10.1101/2022.11.08.515579

**Authors:** Guangzheng Weng, Junil Kim, Kedar Nath Natarajan, Kyoung-Jae Won

## Abstract

Gene regulatory network inference from single-cell RNA sequencing (scRNAseq) datasets has an incredible potential to discover new regulatory rules. However, current computational inference methods often suffer from excessive predictions as existing strategies fail to remove indirect or false predictions. Here, we report a new algorithm single-cell multivariate Transfer Entropy, ‘scmTE’, that generates interpretable regulatory networks with reduced indirect and false predictions. By utilizing multivariate transfer entropy, scmTE accounts for gene-to-gene interdependence when quantifying regulatory relationships. Benchmarking against other methods using synthetic data manifested that scmTE is the unique algorithm that did not produce a hair-ball structure (due to too many predictions) and recapitulated known ground-truth relationships with high accuracy. *In silico* knockdown experiments shows that scmTE assigns higher scores for specific interactions important for differentiation processes. We apply scmTE to T-cell differentiation, myelopoiesis and pancreatic development and identified known and novel regulatory interactions. scmTE provides a robust approach to infer interpretable networks by effectively removing unwanted indirect relationships.

## Introduction

Identifying and understanding gene regulatory rules is crucial to understand various biological processes including cell growth, development and differentiation. Numerous computational approaches have been developed to reconstruct gene regulatory networks (GRNs) from bulk-cell assays including microarrays and RNA sequencing (RNAseq)(Basso, et al., 2005; Hartemink, 2005). However, GRN inference methods from bulk approaches have two major pitfalls. First, the input sample size is extremely low compared with the number of genes (i.e. features assessed). Second, the inference approaches predict too many relationships with a high false positive rate (Nguyen, et al., 2021). The advent of single cell RNA sequencing (scRNA-seq) has enabled simultaneous profiling of transcriptome-wide expression levels across millions of cells, providing an increased sample size for GRN reconstruction (Kim, et al., 2021; Matsumoto, et al., 2017; Pratapa, et al., 2020; Specht and Li, 2017). However, current methods to infer GRNs still suffer from excessive predictions and indirect relationships due to high number of false positives. Algorithmic development of new methods is required that effectively remove false predictions and generate interpretable networks.

Many approaches have been developed for network inference using bulk-RNAseq data using ensembles of regression trees(Huynh-Thu, et al., 2010) or repeated subsampling followed by model training(Morgan, et al., 2020). TRaCE+ utilizes ensembled expression profiles from various knock-out (KO) experiments to define the upper and the lower bound of the networks(Ud-Dean, et al., 2016) and requires multiple genes and multiple linear regression (MLR) to avoid indirect relationships (Salleh, et al., 2017). Even with their efforts, above approaches could not effectively remove indirect relationships.

scRNAseq enables to use dynamic changes of gene expression along the inferred cell-transition or trajectory for inferring GRNs. The GRN inference methods developed using scRNA-seq data focus on investigating relationships between gene pairs followed by an additional measure to remove indirect relationships (Nguyen, et al., 2021; Qiu, et al., 2020). For instance, data processing inequality (DPI), developed previously for GRN inference using bulk RNAseq (Margolin, et al., 2006), was applied to trim the indirect relationships (Kim, et al., 2021). On the other hand, SCODE (Matsumoto, et al., 2017) tried to model the dynamics of a set of genes with ordinary differential equations (ODEs). However, these *post hoc* approaches as well as ODE based approach did not remove the unwanted indirect relationships effectively and often produced hairball network structure due to too many predictions. Therefore, accounting for gene inter-dependencies during the GRN inference step could remove unwanted indirect relationships. However, it is not easy to identify true relationships while many genes are interdependent with each other. Also, it will be computationally exhaustive to consider all relationships among genes. However, there are few approaches trying to identify true relationships while genes are interdependent with each other, because of exhaustive computation consuming. A method that can infer direct regulatory relationship from interactive relationships as well as reduce computation cost is required.

Multivariate transfer entropy (mTE) is an algorithm to obtain causal relationships among multiple variables(Mao and Shang, 2017). Compared with transfer entropy (TE) which calculates causal relationships between two variables, mTE can potentially remove the indirect influence(García-Medina and Hernández C, 2020), and has been applied to a variety of studies including functional magnetic resonance imaging (fMRI) to identify the underlying directed information structure between brain regions considering multivariate sources(Lizier, et al., 2011; Novelli, et al., 2019).

We hypothesize that mTE has potential to remove indirect relationships during the network inference and developed scmTE. scmTE employes mTE to calculate causal relationships between a gene pair while considering other sources (genes) that can influence the target gene expression. To prune the genes whose influences towards the target gene are negligible, scmTE adopts an iterative greedy algorithm(Faes, et al., 2011; Lizier and Rubinov, 2012), and the calculated mTE value (between two genes) becomes the weight of the inferred networks.

To demonstrate the performance of scmTE, we benchmarked scmTE against other methods using simulated scRNA-seq datasets with various developmental trajectories. scmTE successfully recapitulated the reference network structures with a remarkable similarity while other approaches often produced a hair-ball-like structure with an excessive number of predictions. Even when limiting the number of predictions, scmTE demonstrated superior performance compared with other inference methods. We find that the interactions with high weights correspond to crucial factors for pseudotime trajectories.

Applying scmTE to T-lymphocyte differentiation, we captured critical regulatory relationships for both naive T-cells and Th2 differentiated cells. scmTE captured key antagonistic transcription factors and their targets essential for monocyte and granulocyte differentiation, which we re-confirm using knockout scRNA-seq data analysis for both transcription factors. Applying scmTE on a pancreatic development dataset(Bastidas-Ponce, et al., 2019), we detected a GRN centered on Neurog3, a key regulatory and clinical factor driving islet cell formation. In summary, scmTE infers GRNs from scRNA-seq data with robust target predictions and captures key regulatory factors effectively.

## Results

### scmTE considers all possible relationships among genes in GRN reconstruction by applying scmTE

scmTE calculates causal relationships from a gene to a target gene while considering other genes that can influence the target. Similar to TE, mTE relies on the dynamic gene expression changes over time *i*.*e*. pseudo-time, the ordered trajectory from scRNAseq (Figure 1A). To reduce the scmTE computing cost, we select genes within each dataset that have the potential to influence the target gene using a greedy algorithm(Faes, et al., 2011). Briefly, the greedy algorithm iteratively collects a set of source candidates that maximizes the conditional mutual information (CMI) to the target genes (Methods), then prune redundant sources by minimum statistic (Figure 1B). Finally, scmTE produces the directed graphs whose weights are the calculated mTE values (Figure 1C). Moreover, the key regulations can be identified with high mTE values. As the potential sources are considered during the network reconstruction, scmTE does not require any *post hoc* algorithm to remove indirect relationships

**Figure 1.**
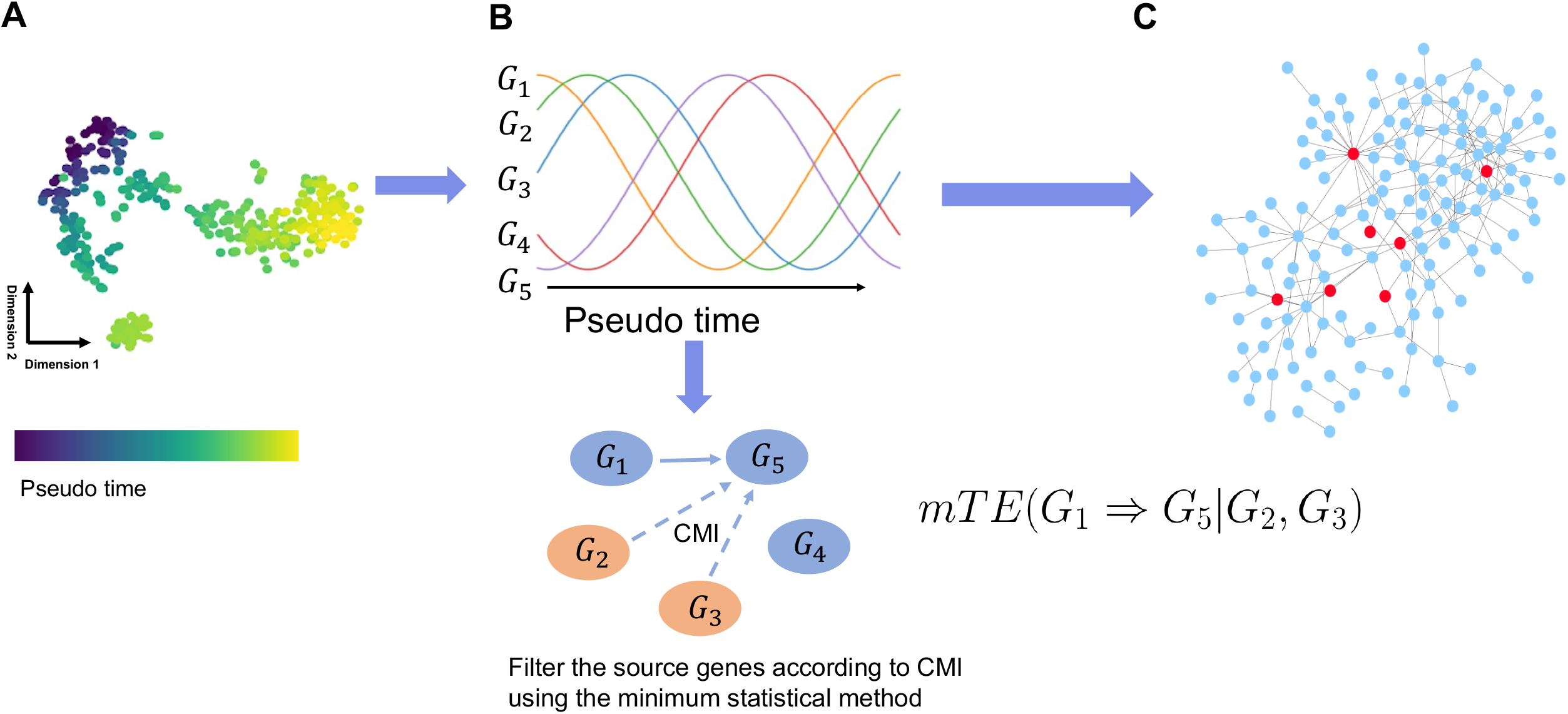
scmTE employs mTE for interpretable GRN by reducing false predictions. A) scmTE uses ordered scRNAseq data from trajectory analysis. B) scmTE identifies the source genes influencing the target genes. mTE is calculated only using the source genes. A greedy algorithm determines that G_2_ and G_3_ as well as G_1_ influence the expression of G_5_ based on CMI. mTE (G_1_ -> G_5_) is calculated while accounting for the past events of G_2_ and G_3_. C) mTE is calculated for all gene pairs to reconstruct the entire GRNs. Red nodes mean key factors with high mTE values.

### Artificial data generation from the networks to assess the performance

The performance evaluation using real scRNAseq datasets is challenging as the true ground truth relationships between genes are known. Therefore, we prepared four network structures (linear, bifurcation, trifurcation and cycle) from simulated scRNAseq datasets using BoolODE(Pratapa, et al., 2020). The simulated data consist of transcriptomes from 2,500 single-cells (Supplementary Figure1).

### Previous GRN inference approaches always produced hairball structures

For benchmarking, we tested the state-of-the-art GRN inference methods including TENET(Kim, et al., 2021), GENIE3(Morgan, et al., 2019), SCODE(Matsumoto, et al., 2017), GRISLI(Aubin-Frankowski and Vert, 2020), GRNBOOST2(Moerman, et al., 2019), GRNBEVM(Sanchez-Castillo, et al., 2018), LEAP(Specht and Li, 2017), SCRIBE(Qiu, et al., 2020) and SINGE(Deshpande, et al., 2021). We included TENET and SCRIBE as they are designed based on the TE algorithm. GENIE3 employed ensembles of regression trees to reduce false predictions(Pratapa, et al., 2020). LEAP implemented a permutation strategy to estimate false-positive connections(Specht and Li, 2017). We also included SCODE and GRISLI as they account for the relationships among multiple genes when reconstructing ODEs from scRNAseq(Aubin-Frankowski and Vert, 2020; Matsumoto, et al., 2017).

We tested the GRN reconstruction algorithms without limiting the number of relationships, as both the number of relationships and the network size are usually unknown. The inferred GRNs across all tested algorithms (GRISLI, GRNBOOST2, GRNVBRM, LEAP, SCRIBE, SINGE) produced a hairball structure, regardless of the datasets and input reference network (Supplementary Figure 1). The observations indicate that existing methods fail to address the issue of indirect relationships or false positives (FPs).

### scmTE can uniquely determine network size and recapitulates the backbone of the reference architecture

We tested scmTE without limiting the size of the networks and assessed its usefulness in removing indirect relationships. Figure 2A shows the single-cell embedding on tSNE, as well as reference network and scmTE prediction; with correctly predicted edges in red and false positives in black. We do not show false negatives (FNs) (when reference edges are unpredicted) for simplicity. The scmTE inferred structure has close similarity with the reference networks (Figure 2A). For instance, the circular backbone for the cycle and the linear structures were almost perfectly inferred by scmTE. The bifurcation and the trifurcation also showed similar structures and most of the inferred relationships were true positives (TFs) (red in Figure 2A). The results by scmTE are striking compared with the results by other predictors (Supplementary Figure 1). Our results show that scmTE can effectively remove the indirect relationships and capture the network topology of underlying scRNAseq data.

**Figure 2.**
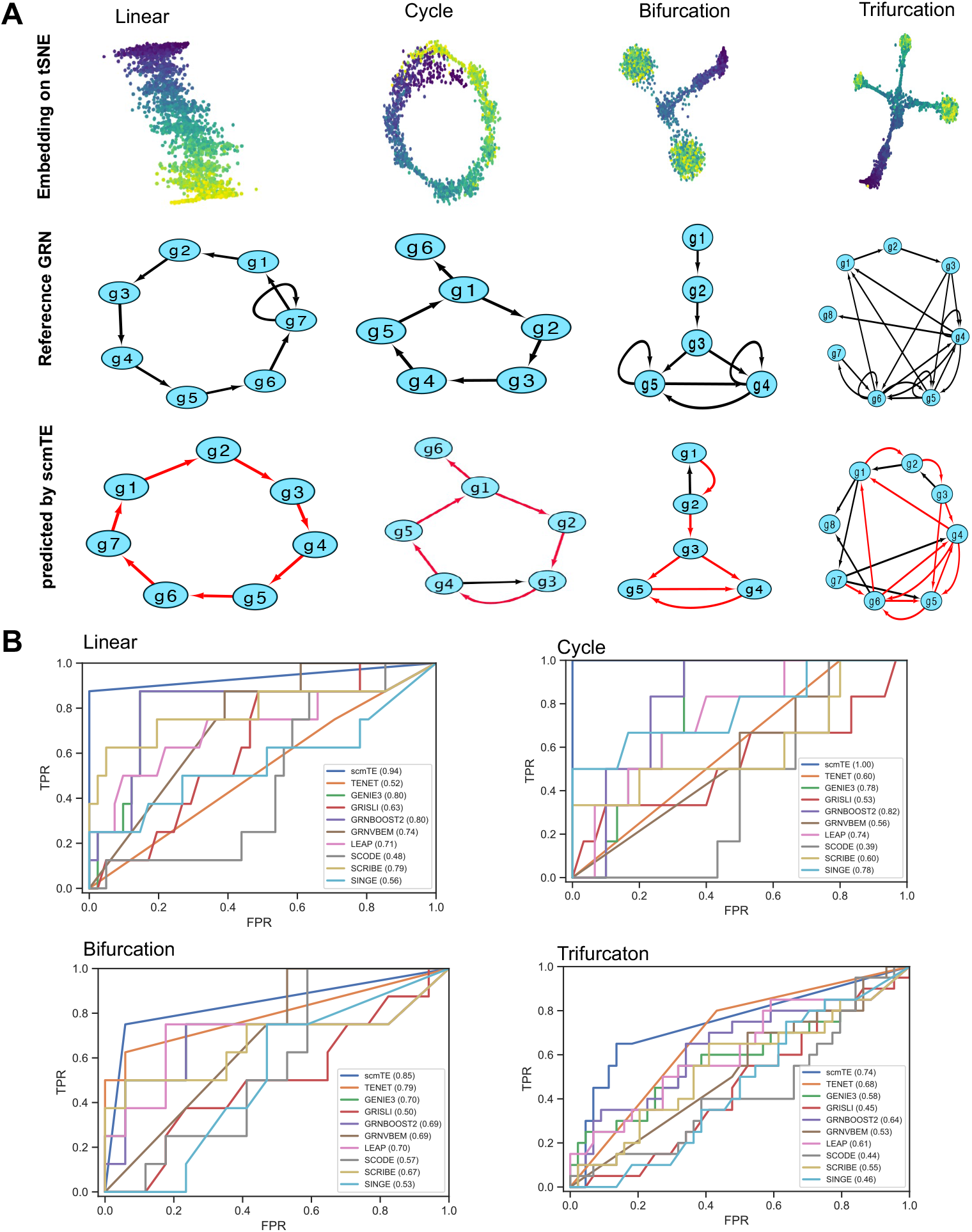
scmTE correctly infers the GRNs from scRNAseq data. A) The inferred GRNs by scmTE for various scRNAseq topology: linear, cycle, bifurcation and trifurcation. The reference GRNs and inferred GRNs are shown. The TP is shown in red and FP is in black. For the predicted GRNs. For simplicity, FN is not shown. B) The comparison of the performance using the 4 trajectories. The area under the receiver operating curve (AUROC) was calculated for all algorithms.

### The top scoring weights by other GRN inference tools still inaccurate

We next assessed whether the top scoring relationships of the inferred GRNs match well with the reference. We used reduced networks composed of top scoring weights (up to the number of the weights in the reference) from the inferred networks (Supplementary Figure2). Compared to the scmTE inferred architecture (Figure 2A), other approaches only show partial overlap between structures (Supplementary Figure2A). For instance, for the linear topology, only scmTE predicted a complete ring structure, while TENET, GRNBOOST2, LEAP and SCRIBE predicted partial structures and other approaches failed to predict underlying target relationships. For the cyclic dataset, scmTE predicted all true-positive connections with only one error (g4->g3) which has the lowest mTE value, while the backbone circular structure was lost in other approaches (Figure 2A). Both SCODE and GRNVBEM failed to provide true-positive connections (Supplementary Figure 2B).

The bifurcation dataset has a root branched into two lineages (g3 to g4 and g5) (Figure 2A). scmTE correctly predicted six edges with 1 inverse edge prediction (g2->g1) (Figure 2A). The underlying edge relationships were lost across GRISL, GRNVEBM, LEAP and SINGE (Supplementary Figure2C). TENET performed second best with five true-positive connections with three false-positive connections (g2->g1, g2->g4 and g3->g2). LEAP also predicted five true-positive connections, but it did not find any relationships associated with g1. GRNBOOST2 and GENIE3 only predicted four true-positive connections. Even though the relationships among genes can be considered in the ODEs, SCODE(Matsumoto, et al., 2017), the ODE based GRN method, did not predict the relationships among genes correctly. For trifurcation dataset, scmTE again showed the highest number of true-positive connections covering all genes (Figure 2A), while other approaches only detected a small portion of the reference GRN (Supplementary Figure2D). In sum, the top scoring relationships of other inference tools miss key links and potentially suffer from indirect relationships.

### scmTE outperforms other GRN predictors

We next assessed the overall performance by investigating the true positive rate (TPR) against the false positive rate (FPR) using the weights or p-values as the cut-off. scmTE showed almost perfect performance for the linear and the cycle structure (area under curve (AUC)>0.9) and outperformed other methods for bifurcation and trifurcation structure (Figure 2B). TE based TENET showed the second-best performance for bifurcation and the trifurcation topology but the performance was similar to random guess (AUC=0.5) for the linear and the cyclic topology. SCRIBE, another TE based approach, was successful for the linear and the bifurcation topology (ROC>0.8) but not for the trifurcation and the cyclic topology. Overall, scmTE showed the best performance regardless of the topological structure.

We further investigated positive predictive value (PPV) while we increase the number of predictions (Supplementary Figure3). The number of interactions from the reference GRNs are shown in the vertical black dotted line, with consistent worsening of PPV as the number of predictions increases. Therefore, it is important to find a suitable cut-off to restrict GRN growth; and scmTE selected the closest number of interactions relative to the reference (Supplementary Figure3). Overall, the number of FPs for scmTE is low and the true positive rate (TPR=TP/(TP+FP)) was over 0.8 for the linear, the bifurcation and the cycle topology and over 0.6 for the trifurcation topology.

### The identified networks re-capitulated the characteristics of the original networks

We further analyzed the weights of the inferred networks by performing *in silico* knockout using the trifurcation model. We removed the edges one by one in an order of the mTE value from the reference. After removing the edge of the network, we re-simulated the scRNAseq and embedded them into 2D t-SNE space (Figure 3A). The knockout of top three connections with high scmTE values: g4->g5, g5->g6 and g5->g4 (Figure 3B) led to seriously deformed topologies from the original trifurcation structure. Compared to the top three connection, the knockout of the three lowest scmTE edge values: g2->g1, g6->g1 and g4->g1(Figure 3B) did not significantly change the cell lineage topology This indicates that scmTE captures functionally important relationships between genes from scRNAseq data.

**Figure 3.**
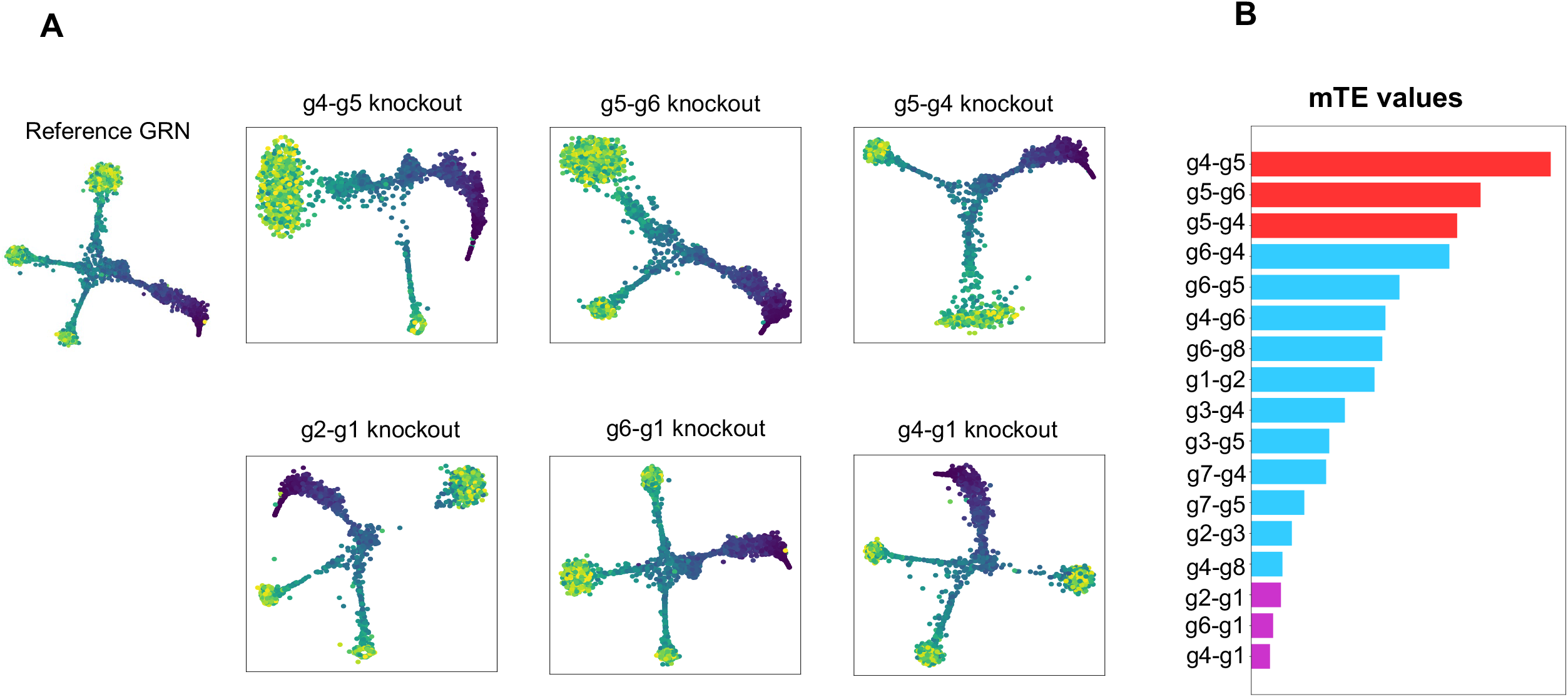
scmTE captures key relationships important for the formation of trajectory. A) In silico knockout of relationships with high mTE values (g4-g5, g5-g6 and g5-g4) leads to failure of trifurcation formation. In silico knockout of relationships with low mTE values (g2-g1, g6-g1,g4-g1) does not influence the building of trajectory. B) The mTE values for all relationships.

### scmTE captures key TFs and novel regulatory relationships during T-cell differentiation

We applied scmTE to the scRNA-seq dataset consisting of naive T cells, and their differentiation to Th2 cells. In addition to key transcription factors (GATA3) and cytokines (IL4, IL5, IL13), splenic differentiation of naïve cells to Th2 cells is a heterogenous process and involves many unknown regulatory relationships(Pramanik, et al., 2018; Radtke, et al.). We performed trajectory analysis using DPT(Haghverdi, et al., 2016) and captured the progression from two naive T-cell states to Th2 cells, and confirmed marker gene expression (Supplementary Figure 4). Besides known regulators such as Il13, we expected many other factors to contribute to the underlying Th2 lymphocyte regulatory network. scmTE successfully identified key regulators expressed at low levels including Il13(Gallo, et al., 2012), Cd5, Cd6 (T-cell activation and proliferation), Cd52 (T cell marker)(Samten, 2013) (Figure 4A). Notably, scmTE captured Il13 as a large hub with several target genes including Cd4, Gata3, Ccr8, and Srpr (Figure 4B). scmTE can contrast direct regulatory relationships important for T-cell activation, proliferation, and differentiation from Th2 activity. These include Ly6a(Malek, et al., 1989), Tnfsrf4(Schreiber, et al., 2012), CD2(Binder, et al., 2020), Tnfsf8(T-cell survival)(Yang, et al., 2020), BCAP31 (Highly expressed T-cell enriched transcript)(Quistgaard, 2021), and Il12rb1(Reeme, et al., 2019). We describe all the regulatory targets and links in Supplementary Table1. In summary, scmTE captures and predicts key regulatory factors important for naive T cell differentiation.

**Figure 4.**
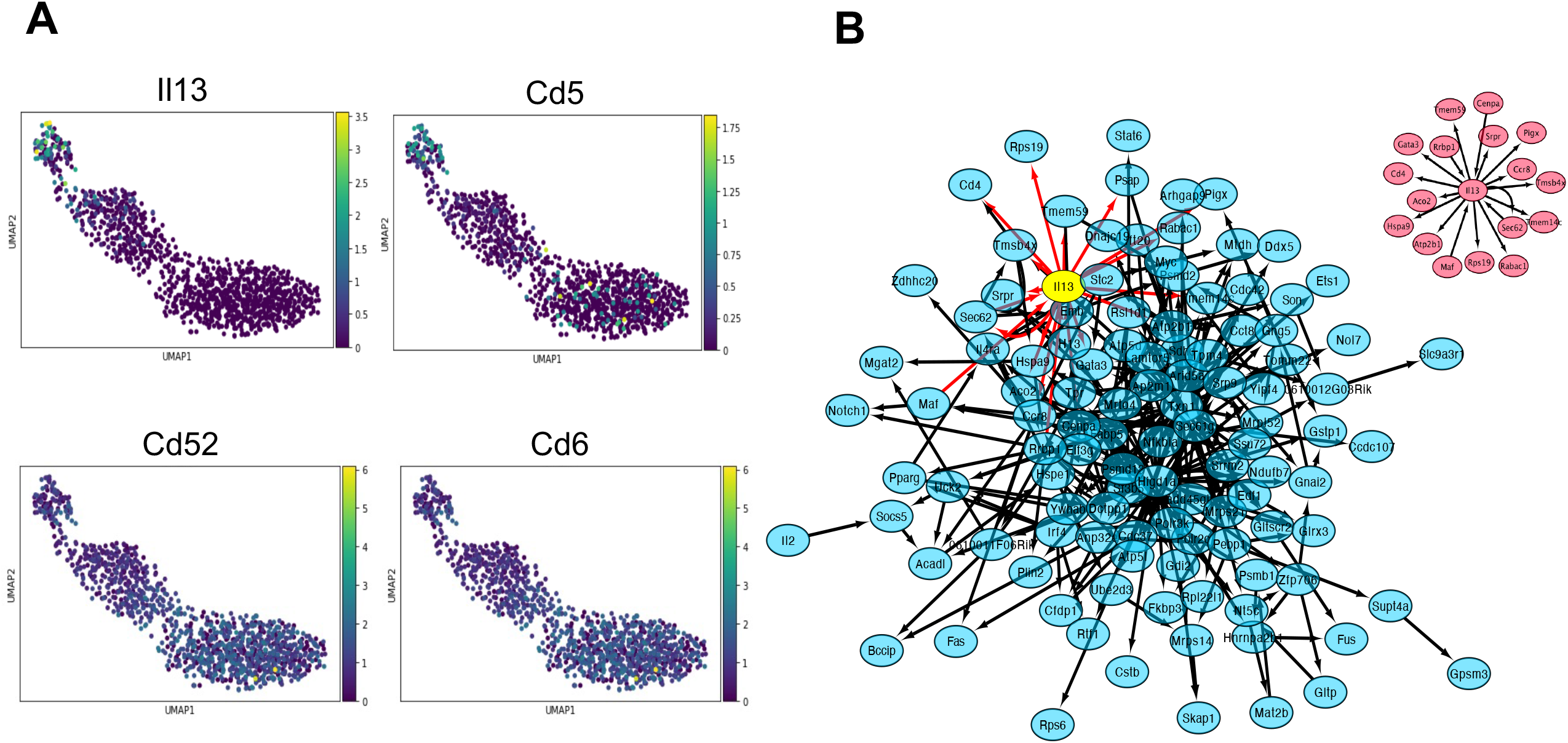
scmTE captures regulatory factors for T cells. A) Expression changes for Il13, Cd5, Cd52 and Cd6. B) The inferred GRNs by scmTE. The network involving the key regulatory factors Il13 is shown in the network. The predicted target genes for Ill3 are separately shown.

### scmTE captures key TFs that resolve mixed-lineage states during myelopoiesis

Myelopoiesis involves a hierarchical cell fate specification of discrete hematopoietic multipotent progenitors into transitory bipotential intermediates that further generate monocytes and granulocytes(Olsson, et al., 2016). During normal development, these primed lineage intermediates concurrently express both stem and specific myeloid progenitors prior to specification. Notably, the deletion of the counteracting TFs traps cells into metastable intermediates that fail to undertake monocytic or granulocytic differentiation programs.

We tested whether scmTE can infer key regulatory myeloid determinant factors by analyzing scRNA-seq data spanning the myeloid differentiation program, beginning from hematopoietic stem cell progenitors (HSCP1, HSCP2), megakaryocytic (Meg), erythrocytic (Eryth), multi-lineage primed cells (Multi-Lin), monocyte-dendritic precursors (MDP) to monocytic (mono), granulocytic (Gran) and myelocytes (Figure 5A). The inferred pseudo-time recapitulated single-cell trajectory through respective cell types (Figure 5B). Notably, scmTE captured the lowly expressed but key antagonistic TFs critical for monocyte (Irf8, Klf4) and granulocyte (Gfi1, Cebpe, Elane, Tmem2, Afap1) specification respectively, alongside broad acting TFs across progenitor cells (Gata2, Meis1, CEBP) (Figure 5C and D). scmTE captured essential TF-target relationships in monocyte (Irf8-Klf4) and across granulocytes (Gfi1-Lgals3, Gfi1-Tmem2, Gfi1-Afap1) (Supplementary Figure5A), capturing the regulatory architecture within discrete cell states.

**Figure 5.**
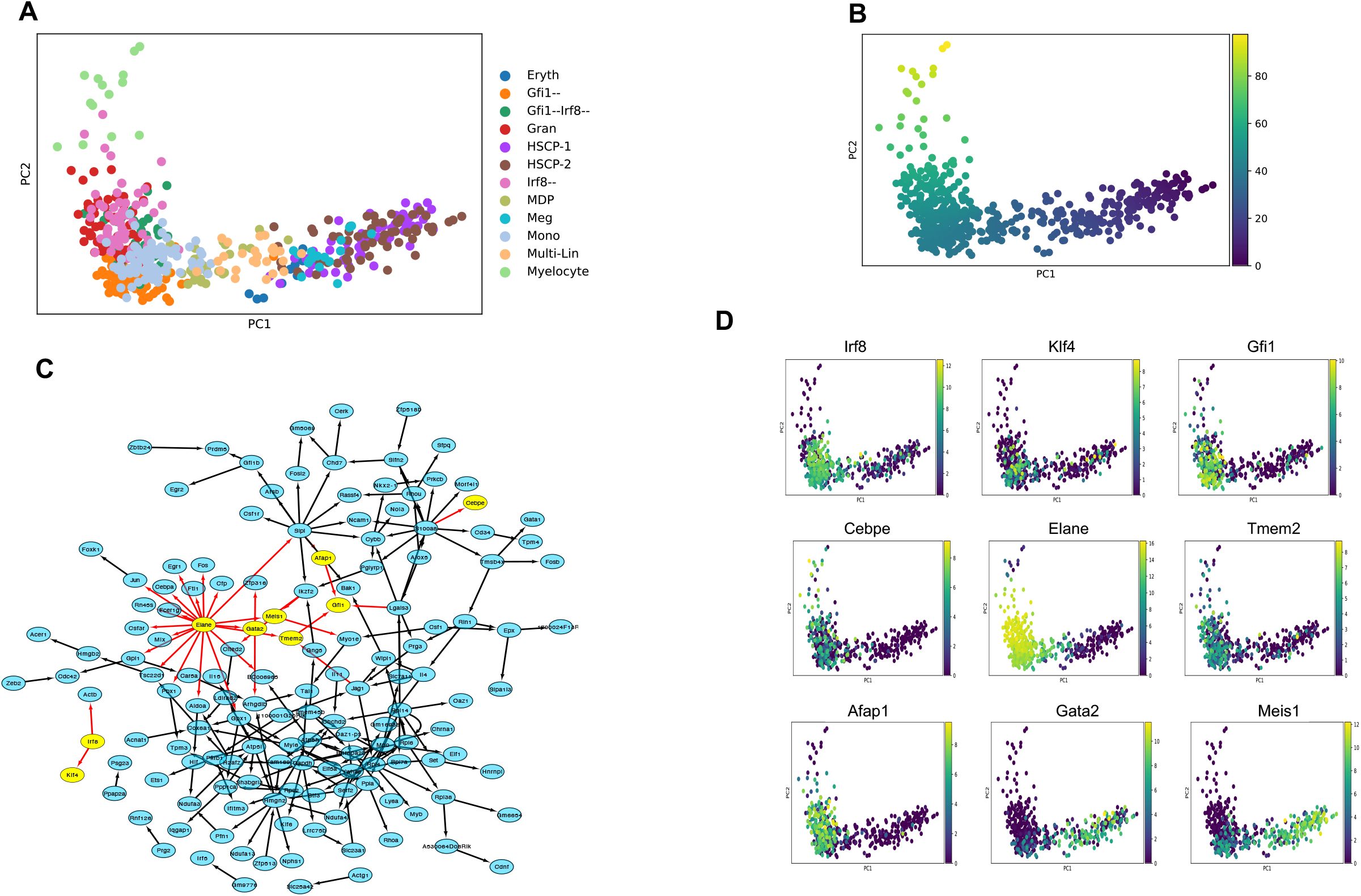
scmTE sensitively captures key TFs that resolve mixed-lineage states during myelopoiesis. A) scRNAseq data are embedded on 2D PCA space. B) The trajectory analysis by Slingshot shows that erythrocytic cells differentiate to myelocyte cells. C) The entire GRN inferred by scmTE. Known key factors are marked by yellow and the inferred relationships are colored by red. D) Expression profile of key factors are shown in the PCA space.

To validate the scmTE predictions, we analyzed knockout cells including Irf8^-/-^, Gfi1^-/-^ and double Irf8^-/-^Gfi^-/-^ scRNA-seq data to assess which TF-targets links were lost and find regulatory relationships in double knockout cells. The knockout impairs protein function through perturbation of specific exons for Gfi1 and Irf8 respectively(Olsson, et al., 2016). We selected the top 100 highly variable features including known targets for scmTE analysis. Globally, the number of TF-target relationships before and upon knockout were similar (201 in Gfi1^-/-^, 198 in Irf8^-/-^, 184 in Irf8^-/-^Gfi1^-/-^; vs 214 total links; Supplementary Table2). However, the number of Gfi1 links decreased upon single and double knockouts (Irf8^-/-^ and Irf8^-/-^Gfi1^-/-^), consistent with a reciprocal increase in Irf8 targets (Supplementary Figure6). We observed that upon Gfi1 perturbation, Gfi1 links were regulated by Irf8 and vice versa in Irf8 perturbation (Supplementary Figure5). Consistently, the Irf8^-/-^ cells had granulocytic specification concomitant with increased expression of Gfi1 (targets Per3, Ets1, Cybb-Gfi1, Il4-Gfi1) and skewed regulatory network (Irf8->Eif5a, Irf8->Rassf4, Irf8->Csf1r; Supplementary Figure5B); while the Gfi1^-/-^ had reciprocal induction of monocytic differentiation (Irf8 & targets Klf4, Xeb2, Irf5, Afap1, B4galt5) and regulatory relationships (Irf8->Cebpa, Irf8->Elane; Supplementary Figure 5C). Notably, applying scmTE to Irf8^-/-^ Gfi1^-/-^ cells failed to predict the known Irf8 and Gfi1 TFs-targets relationships, instead captured a highly perturbed network of antagonistic relationships (Irf8->Cebpa,Irf8->Elane,Irf8->Csf1r, Slpi->Gfi1, Gfi1->Chd7, Irf8->Tpm4, Rn45s->Irf8, Irf8->Rhoa)(Supplementary Figure 5D), validating the double negative cells revert to a metastable population without a defined regulatory program.

### scmTE finds crucial TF for pancreatic development

Finally, we applied scmTE to endocrine development in the pancreas, which comprises two main cell fates: α and β. Endocrine cells originate from the endocrine progenitors, marked by a key TF Neurog3. A previous study proposed that Neurog3 can start endocrine development by regulating a set of TFs that are related to the proliferation and maturation of endocrine cells (Napolitano et al. 2015). Islet cells cannot be formed in mice lacing Neurog3, leading to diabetes(Schreiber, et al., 2021).

The pancreatic datasets are collected from mouse embryonic day 15.5. After preprocessing and analysis, 2531 cells are embedded into reduce UMAP latent space (Figure 6A), where the pseudo time shows a clear cell development trajectory from Ngn3 low EP to α, β, δ, and ε cells (Figure 6B). scmTE captured a small GRN centered on Neurog3, with 15 targets (Figure 6C). Several targets (Insm1, Irx1, Irx2, Neurod2, Pax4) have been shown to play key roles in embryonic development. Pax4 is an important regulator for β cells construction. Experiments show that mice lacking Pax4 die after a few days because they lack mature β cells(Napolitano, et al., 2015; Schreiber, et al., 2021). Insm1 maintains mature β-cell functions(Jia, et al., 2015). Irx1 and Irx2 were not expressed in Neurog3 mutant embryos(Jia, et al., 2015; Petri, et al., 2006). Moreover, scmTE found a regulatory interaction between Neurog3 and Arx, which is still poorly understood. However, previous experiments demonstrated that Arx expressed in normal mouse pancreas, but not in Neurog3 null mice(Collombat, et al., 2003). In summary, Neurog3, a crucial factor that has been widely shown to play an important role in islet cell development, was successfully detected with many targets using scmTE.

**Figure 6.**
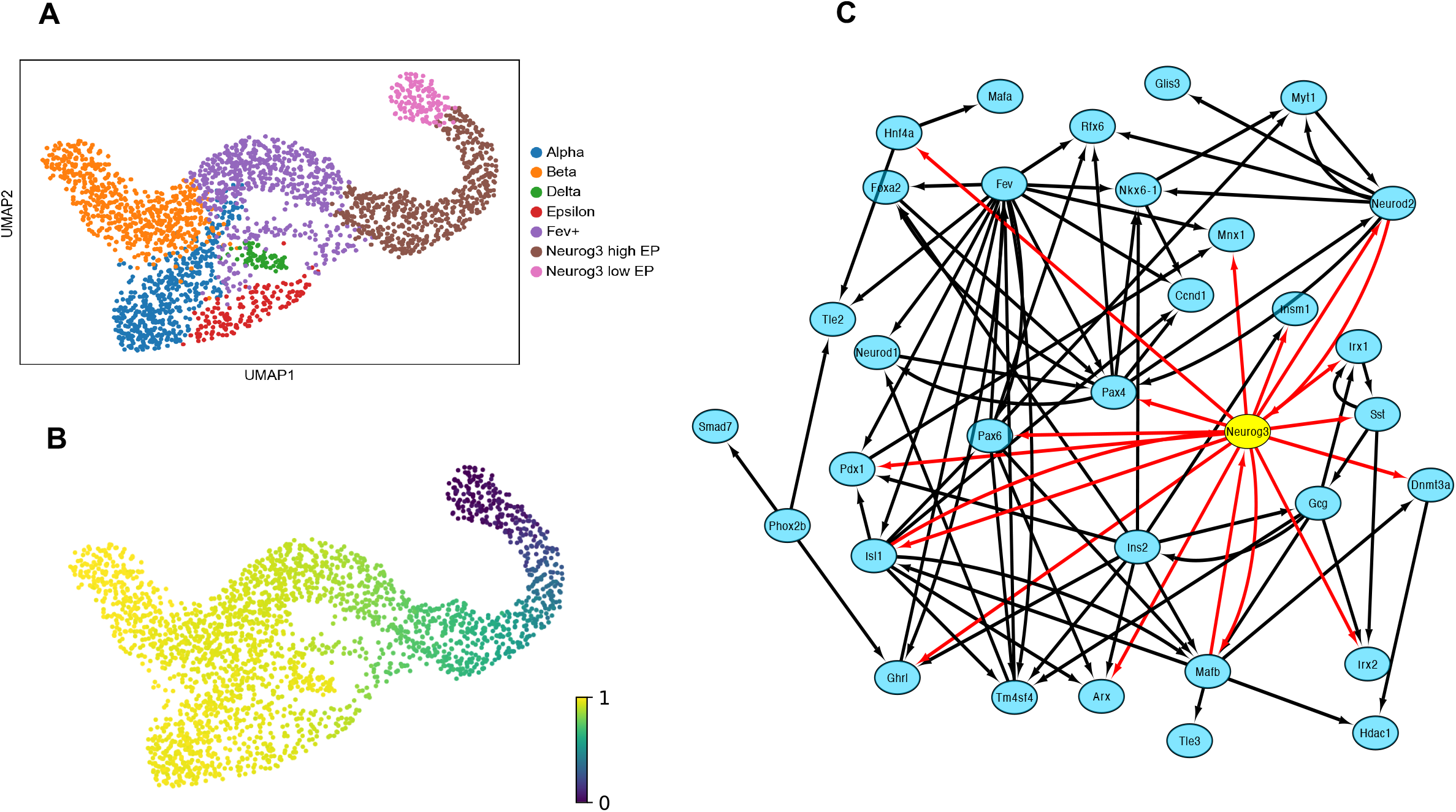
scmTE captures a GRN centered on Neurog3 from the scRNAseq data for pancreas development. A) 2,531 cells from endocrine development in the pancreas embedded on 2D UMAP. Cell types including beta, alpha and Neurog3+ EP cells are shown. B) A pseudo time showed the trajectory from Neurog3+ cells to four cell fates (alpha, beta, delta and epsilon). C) scmTE reconstructs a GRN, where Neurog3 forms a dense network, suggesting ath Neurog3 is a key player during endocrine cell differentiation.

## DISCUSSION

The computational reconstruction of GRNs provides an avenue to infer new gene regulatory rules from expression data(Hartemink, 2005). While a large number of algorithms have been developed for bulk and scRNAseq(Wanqi, 2021), these methods still lack accuracy and interpretability as they suffer from predicting indirect relationships. As shown across our benchmarks, most existing methods predict excessive relationships and lack measures to remove indirect relationships (Supplementary Figure1). Although a widely used approach is to apply a strict threshold to the initially inferred networks thereby ignoring weak interactions; such methods lead to inaccurate prediction of strong interactions (Supplementary Figure2).

Here, we present scmTE, an GRN inference approach that utilizes mTE to produce interpretable GRN with fewer but robust predictions from scRNAseq data. scmTE quantifies the transfer of information while accounting for co-dependencies among genes. Benchmarking using synthetic datasets with known reference GRN architecture showed remarkable advantages of scmTE against other approaches. Without any threshold, scmTE determined the size of the networks (Figure 2). This made scmTE avoid producing non-interpretable hair-ball structures. Moreover, the prediction by scmTE was quite accurate compared with other competitors even when we consider only top scoring interactions (Figure 2 and Supplementary Figure 2). We further confirmed that scmTE puts more scores on the interactions important for the scRNAseq trajectory (Figure 3). In sum, scmTE is an approach to produce robust and compact GRNs without any arbitrary cut-off, a parameter often required by other methods.

Previous GRN reconstruction approaches often used *post hoc* approaches to remove indirect relationships (Pratapa, et al., 2020). The *post hoc* approaches, however, cannot deal with the relationships of all genes. On the other hand, scmTE used the co-dependency information among genes when scoring the relationships. This strategy, compared with previous TE-based approaches (Kim, et al., 2021; Qiu, et al., 2020), effectively removed false positive predictions. SCODE (Matsumoto, et al., 2017) could handle the co-dependency when optimizing the parameters of the ODEs. However, it may be difficult to calculate all co-dependencies in the ODE.

We apply scmTE to real scRNA-seq datasets spanning T cell differentiation, myelopoiesis and pancreatic differentiation, confirming known and identifying novel TF-target relationships. Notably, scmTE robustly captures important transcription factors-target relationships across both discrete and continuous cell states (Figure 4 and 5). In conclusion, scmTE is a robust and interpretable GRN inference approach for scRNA-seq datasets and can be broadly applied to identify causal TF-target relationships.

## Methods

The traditional transfer entropy quantifies the reduction of uncertainty of a target process *Y* from a source process *X*. In a multivariate context, the mTE is defined as the information that a source *X* provides for a target *Y* conditioning all the other relevant sources in the network. Unlike the ordinary TE, scmTE requires a set of relevant sources for each target. For particularly small networks, a brute force approach tests all nodes in the network to find suitable sources. However, it is too time-consuming and memory-consuming to apply this on normal-sized networks. Hence, Lizier(Lizier and Rubinov, 2012) and Faes(Faes, et al., 2011) came up with a greedy and iterative algorithm for mTE quantification. The idea of this algorithm is to select all relevant sources of a given target by picking up the variables(which are defined from the pasts of the sources) for each source that contributes the maximum CMI to the target and the selected variables form a multivariate, non-uniform embedding of source genes(Faes, et al., 2011). This algorithm can build a set of sources and remove redundancies for every target.

We define a set of variables: *Z* is the relevant sources, *C* is the candidate set of past variables by providing the maximum lag parameter for the target and sources. This parameter picks up the number of past variables in target and sources for estimating *Z. Y* is denoted as the target and *X* is denoted as the source.

1. Initialize *Z* as an empty set and collect the candidate sets of *Y*’s past (*C*_*Y*_) and *X*’s past (*C*_*X*_)
2. Select variables from *C*_*Y*_: A set of candidate variables c Є C is defined from the pasts of the sources X. For each candidate in *C*_*Y*_, measure the mutual information from *c* to *Y*_*n*_ as *I* (*c*; *Y*_*n*_/ *Z*) Pick up the candidate with maximum mutual information, *c*\*, and perform a significance test, if significant, add *c*\* to *Z* and remove it from *C*_*Y*_. Terminate when *I* (*c*; *Y*_*n*_/ *Z*) is unsignificant or *C*_*Y*_ is empty.
3. Select variables from *C*_*X*_: Collect variables for *C*_*X*_ following the procedure similar to step 2
4. Prune *Z* : Test and remove redundant variables in *Z* via minimum statics. The conditional mutual information between variable in *Z* and the current value of *Y*_*n*_, conditional on all other variables in *Z*.
5. Perform statistical tests on the final set *Z* : Employ an omnibus test to test the information transfer from all source variables to the target: *I(Z*_*X*_ ; *Y*_*n*_*/Z*_*y*_)

Finally, the mTE between a single source *X*_*i*_ and target *Y* is measured from all past variables in *Z* for a source *X*_*i*_: TE(*X*_*i*_ ; *Y*_*n*_*/Z\X*_*i*_).

scmTE incorporates the mTE measurement algorithm and takes ordered scRNAseq dataset as input. Using pseudo time, scmTE estimates the casual relationship for potential regulators and targets.

### Simulation data generation

Artificial datasets were simulated by BoolODE covering four trajectory topologies: linear, bifurcation, trifurcation and cycle. All simulations were generated using default parameter.

### Data generation and processing

### Pancreatic dataset

Pancreatic dataset was collected from mouse embryo(Bastidas-Ponce, et al., 2019). Preprocessing and pseudo time analysis were implemented by standard pipeline from scvelo(Bergen, et al., 2020).

### Splenic immune cells and single-cell analysis

Naive T cells were isolated from mouse spleens followed by polarization and early differentiation into Th2 subtype, following a 3.5 day protocol, as described in(Proserpio, et al., 2016). Naive and activated Th2 cells were processed and sequenced through 10x chromium single-cell transcriptomics protocol following manufacturer’s instructions.

## Data deposit

https://github.com/wgzgithub/scmTE_data_deposite

## Code available

https://github.com/wgzgithub/scmte_v1

## Acknowledgements

This work was supported by Independent Research Fund Denmark [0135-00243B] to K.J.W. The research in KNN lab is supported by Villum Young Investigator grant (VYI#00025397) and Novo Nordisk grants (#NNF18OC0052874).

## References

Aubin-Frankowski, P.-C. and Vert, J.-P. Gene regulation inference from single-cell RNA-seq data with linear differential equations and velocity inference. Bioinformatics 2020;36(18):4774–4780.

Basso, K., et al. Reverse engineering of regulatory networks in human B cells. Nature genetics 2005;37(4):382–390.

Bastidas-Ponce, A., et al. Comprehensive single cell mRNA profiling reveals a detailed roadmap for pancreatic endocrinogenesis. Development 2019;146(12):dev173849.

Bergen, V., et al. Generalizing RNA velocity to transient cell states through dynamical modeling. Nat. Biotechnol. 2020;38(12):1408–1414.

Binder, C., et al. CD2 Immunobiology. Front. Immunol. 2020;11:1090.

Collombat, P., et al. Opposing actions of Arx and Pax4 in endocrine pancreas development. Genes Dev. 2003;17(20):2591–2603.

Deshpande, A., et al. Network inference with granger causality ensembles on single-cell transcriptomic data. BioRxiv 2021:534834.

Faes, L., Nollo, G. and Porta, A. Information-based detection of nonlinear Granger causality in multivariate processes via a nonuniform embedding technique. Physical Review E 2011;83(5):051112.

Gallo, E., Katzman, S. and Villarino, A.V. IL-13-producing Th1 and Th17 cells characterize adaptive responses to both self and foreign antigens. Eur. J. Immunol. 2012;42(9):2322–2328.

García-Medina, A. and Hernández C, J.B. Network analysis of multivariate transfer entropy of cryptocurrencies in times of turbulence. Entropy 2020;22(7):760.

Haghverdi, L., et al. Diffusion pseudotime robustly reconstructs lineage branching. Nat. Methods 2016;13(10):845–848.

Hartemink, A.J. Reverse engineering gene regulatory networks. Nature biotechnology 2005;23(5):554–555.

Huynh-Thu, V.A., et al. Inferring Regulatory Networks from Expression Data Using Tree-Based Methods. PLoS ONE 2010;5(9):e12776.

Jia, S., et al. Insm1 cooperates with Neurod1 and Foxa2 to maintain mature pancreatic β-cell function. EMBO J. 2015;34(10):1417–1433.

Kim, J., et al. TENET: gene network reconstruction using transfer entropy reveals key regulatory factors from single cell transcriptomic data. Nucleic Acids Res. 2021;49(1):e1.

Kim, J., et al. TENET: gene network reconstruction using transfer entropy reveals key regulatory factors from single cell transcriptomic data. Nucleic acids research 2021;49(1):e1–e1.

Lizier, J. and Rubinov, M. Multivariate Construction of Effective Computational Networks from Observational Data. 2012.

Lizier, J. and Rubinov, M. Multivariate construction of effective computational networks from observational data. 2012.

Lizier, J.T., et al. Multivariate information-theoretic measures reveal directed information structure and task relevant changes in fMRI connectivity. J. Comput. Neurosci. 2011;30(1):85–107.

Malek, T.R., Shevach, E.M. and Danis, K.M. Activation of T lymphocytes through the Ly-6 pathway is defective in A strain mice. J. Immunol. 1989;143(2):439–445.

Mao, X. and Shang, P. Transfer entropy between multivariate time series. Communications in Nonlinear Science and Numerical Simulation 2017;47:338–347.

Margolin, A.A., et al. ARACNE: an algorithm for the reconstruction of gene regulatory networks in a mammalian cellular context. In, BMC bioinformatics. Springer; 2006. p. 1–15.

Matsumoto, H., et al. SCODE: an efficient regulatory network inference algorithm from single-cell RNA-Seq during differentiation. Bioinformatics 2017;33(15):2314–2321.

Moerman, T., et al. GRNBoost2 and Arboreto: efficient and scalable inference of gene regulatory networks. Bioinformatics 2019;35(12):2159–2161.

Morgan, D., et al. Perturbation-based gene regulatory network inference to unravel oncogenic mechanisms. Scientific reports 2020;10(1):1–12.

Morgan, D., et al. A generalized framework for controlling FDR in gene regulatory network inference. Bioinformatics 2019;35(6):1026–1032.

Napolitano, T., et al. Pax4 acts as a key player in pancreas development and plasticity. Semin. Cell Dev. Biol. 2015;44:107–114.

Nguyen, H., et al. A comprehensive survey of regulatory network inference methods using single cell RNA sequencing data. Briefings in bioinformatics 2021;22(3):bbaa190.

Novelli, L., et al. Large-scale directed network inference with multivariate transfer entropy and hierarchical statistical testing. Netw Neurosci 2019;3(3):827–847.

Olsson, A., et al. Single-cell analysis of mixed-lineage states leading to a binary cell fate choice. Nature 2016;537(7622):698–702.

Petri, A., et al. The effect of neurogenin3 deficiency on pancreatic gene expression in embryonic mice. J. Mol. Endocrinol. 2006;37(2):301–316.

Pramanik, J., et al. Genome-wide analyses reveal the IRE1a-XBP1 pathway promotes T helper cell differentiation by resolving secretory stress and accelerating proliferation. Genome Medicine 2018;10(1).

Pratapa, A., et al. Benchmarking algorithms for gene regulatory network inference from single-cell transcriptomic data. Nature methods 2020;17(2):147–154.

Proserpio, V., et al. Single-cell analysis of CD4+ T-cell differentiation reveals three major cell states and progressive acceleration of proliferation. Genome Biol. 2016;17:103.

Qiu, X., et al. Inferring causal gene regulatory networks from coupled single-cell expression dynamics using scribe. Cell systems 2020;10(3):265-274. e211.

Quistgaard, E.M. BAP31: Physiological functions and roles in disease. Biochimie 2021;186:105–129.

Radtke, D., et al. Th2 single-cell heterogeneity and clonal interorgan distribution in helminth-infected mice.

Reeme, A.E., et al. Human IL12RB1 expression is allele-biased and produces a novel IL12 response regulator. Genes Immun. 2019;20(3):181–197.

Salleh, F.H.M., Zainudin, S. and Arif, S.M. Multiple linear regression for reconstruction of gene regulatory networks in solving cascade error problems. Advances in bioinformatics 2017;2017.

Samten, B. CD52 as both a marker and an effector molecule of T cells with regulatory action: Identification of novel regulatory T cells. Cell. Mol. Immunol. 2013;10(6):456–458.

Sanchez-Castillo, M., et al. A Bayesian framework for the inference of gene regulatory networks from time and pseudo-time series data. Bioinformatics 2018;34(6):964–970.

Schreiber, T.H., et al. T cell costimulation by TNFR superfamily (TNFRSF)4 and TNFRSF25 in the context of vaccination. J. Immunol. 2012;189(7):3311–3318.

Schreiber, V., et al. Extensive NEUROG3 occupancy in the human pancreatic endocrine gene regulatory network. Mol Metab 2021;53:101313.

Specht, A.T. and Li, J. LEAP: constructing gene co-expression networks for single-cell RNA-sequencing data using pseudotime ordering. Bioinformatics 2017;33(5):764–766.

Ud-Dean, S.M., et al. TRaCE+: ensemble inference of gene regulatory networks from transcriptional expression profiles of gene knock-out experiments. BMC bioinformatics 2016;17(1):1–14.

Wanqi, Z. The Comparison between Bulk RNA-seq and Ssingle-cell RNA-seq. Journal of Physics: Conference Series 2021;1893(1):012014.

Yang, X., et al. Age-Related Gene Alteration in Naïve and Memory T cells Using Precise Age-Tracking Model. Front Cell Dev Biol 2020;8:624380.

