## Supplementary figures and images for "scmTE: multivariate transfer entropy builds interpretable compact gene regulatory networks by reducing false predictions"

### Supplementary Figure1

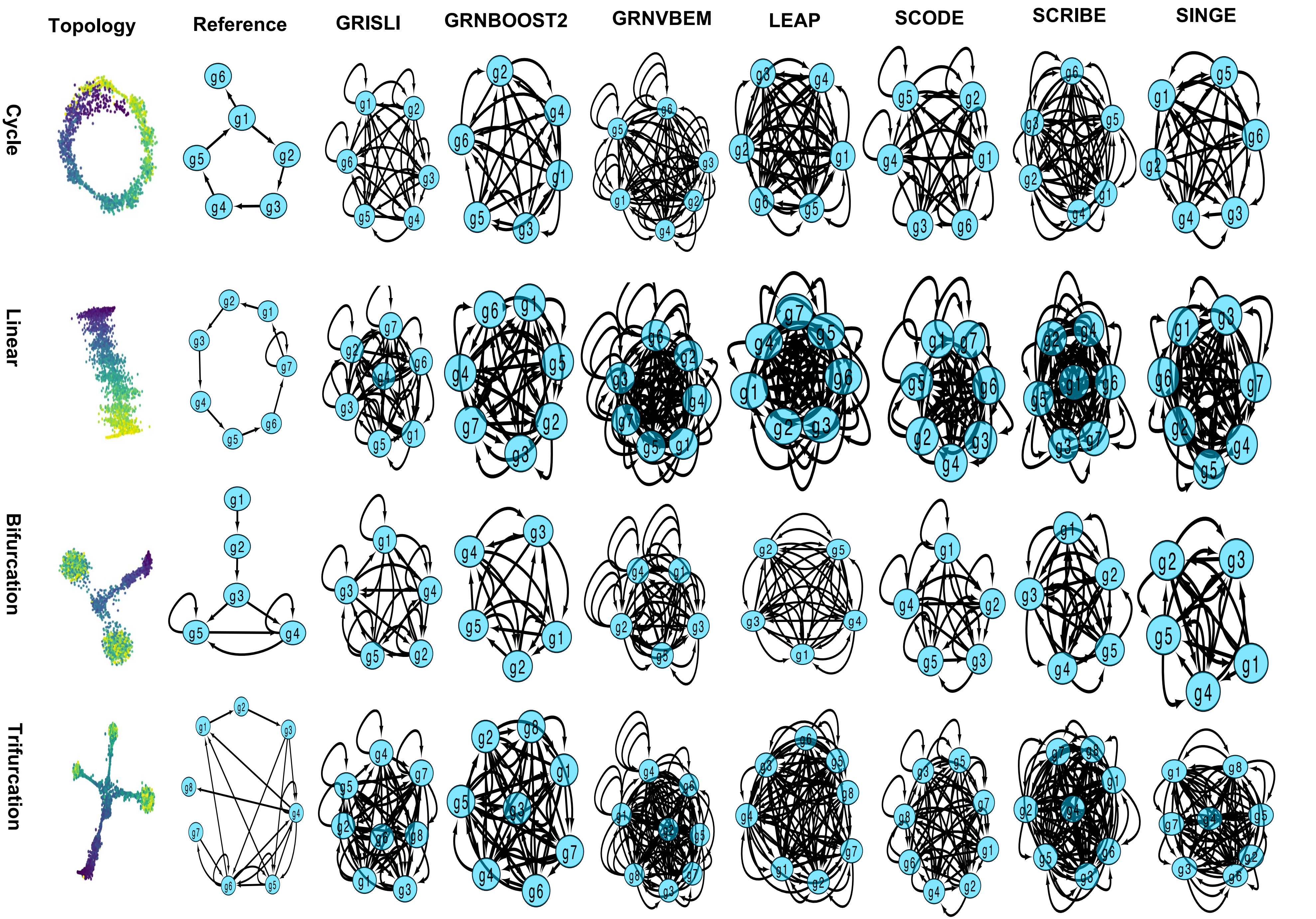

### Supplementary Figure2

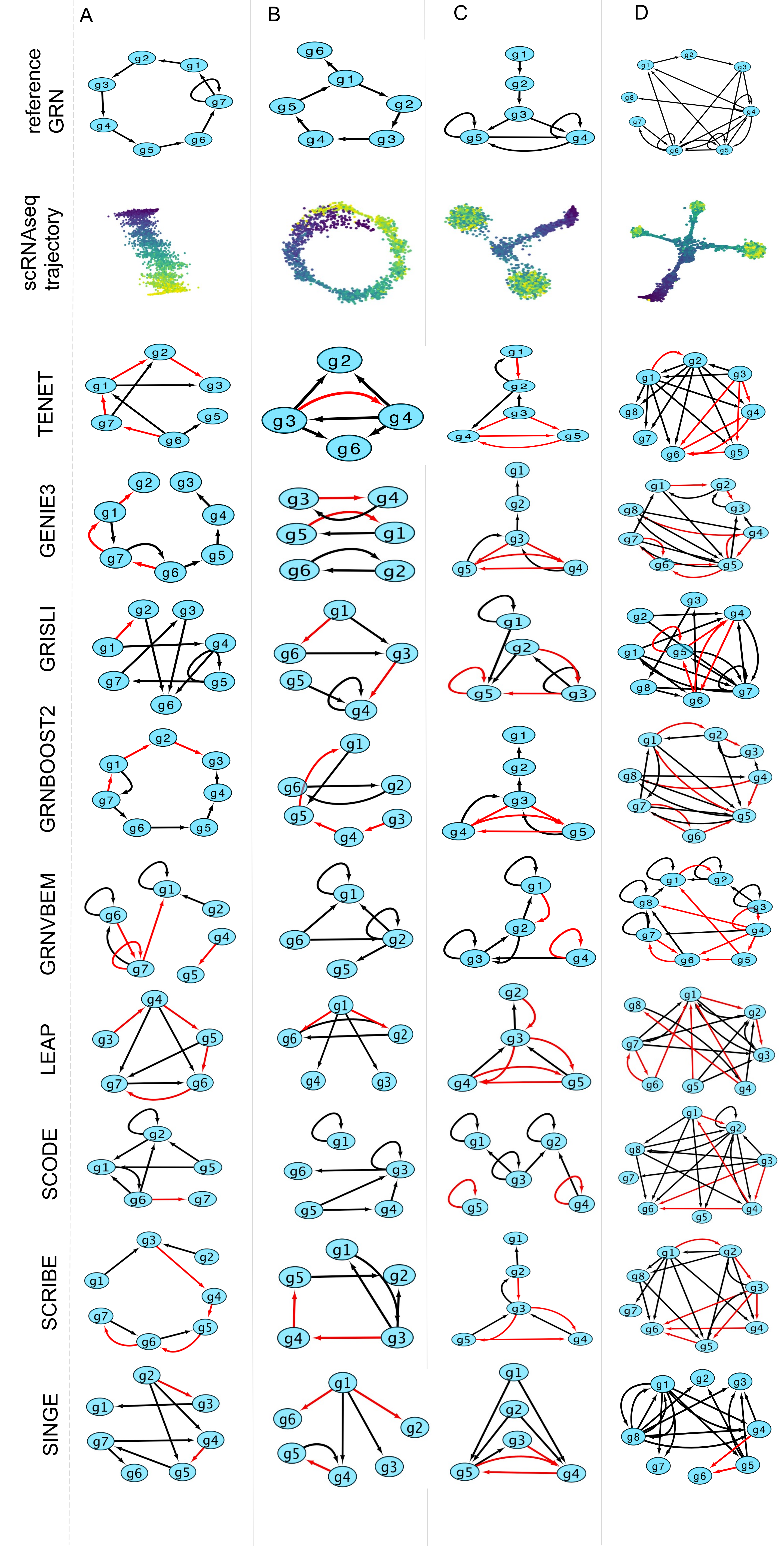

### Supplementary Figure3

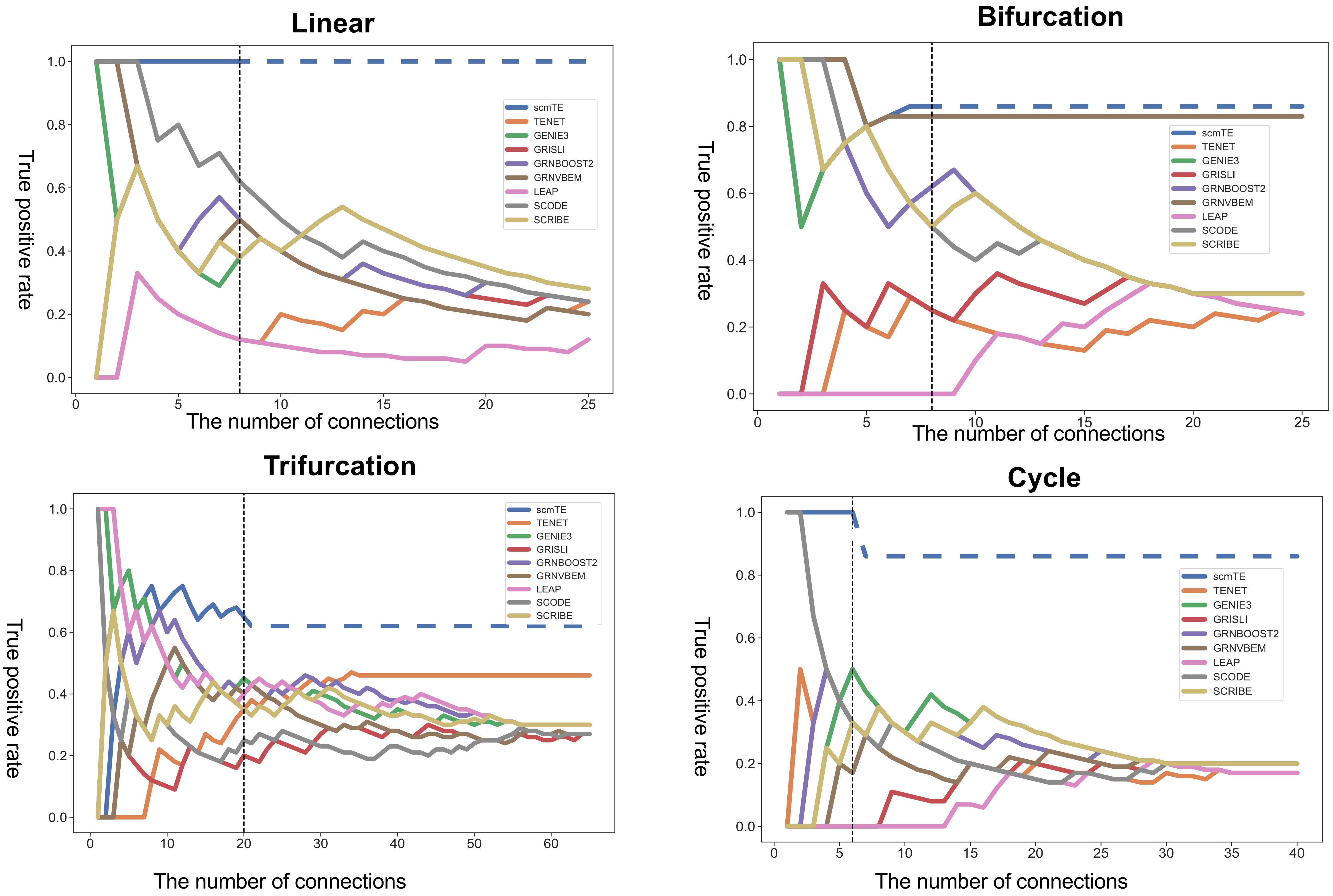

### Supplementary Figure4

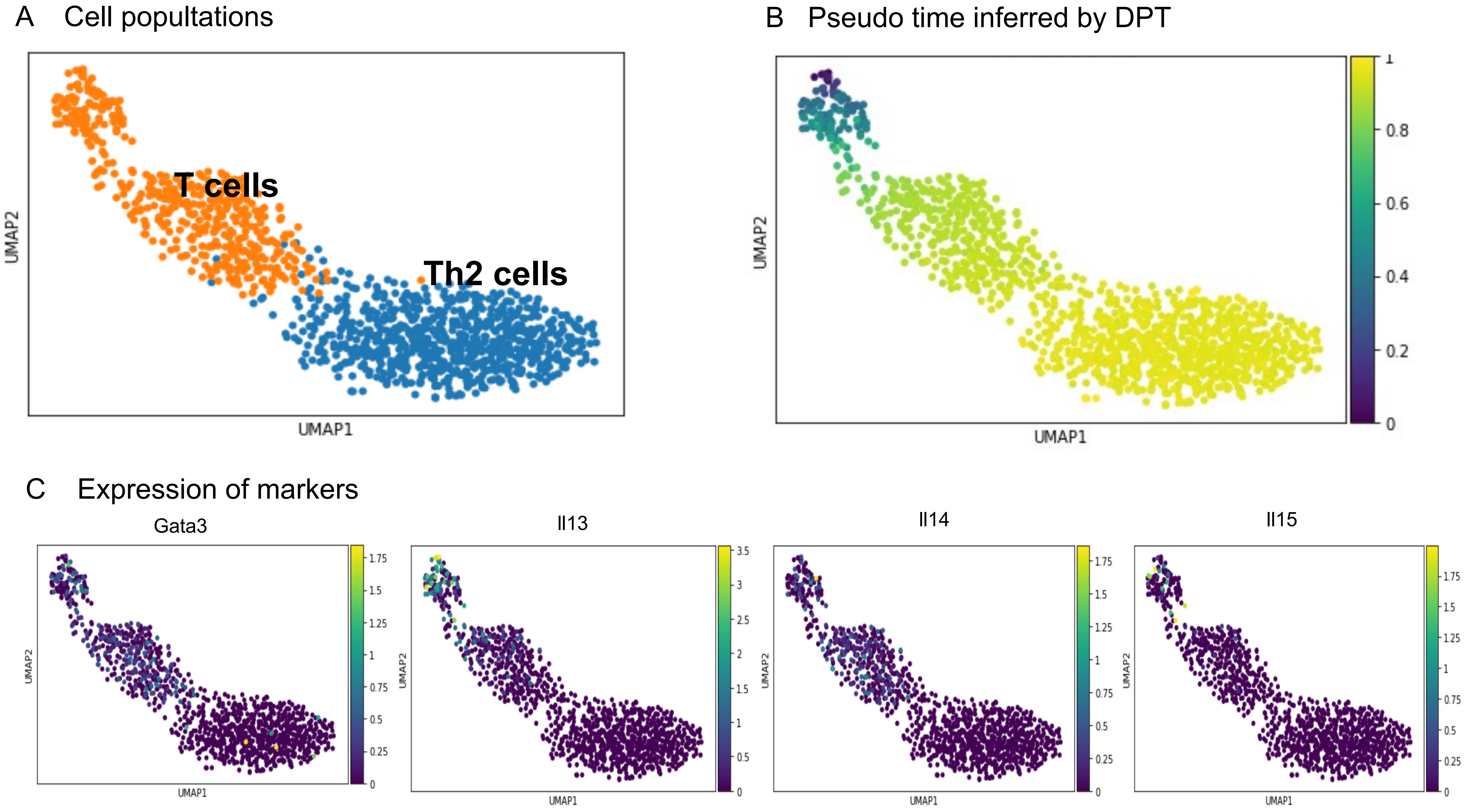

### Supplementary Figure5

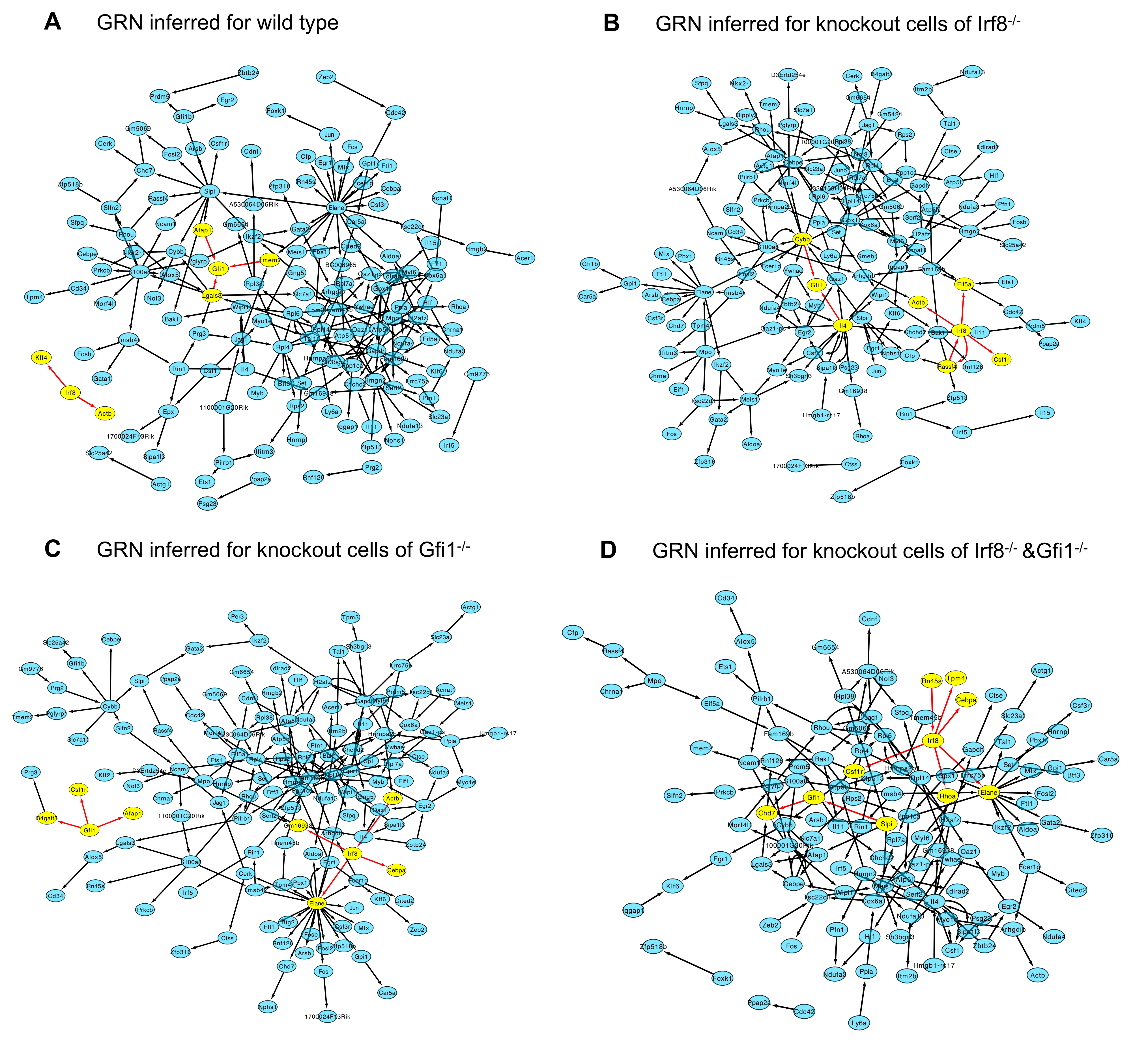

### Supplementary Figure6

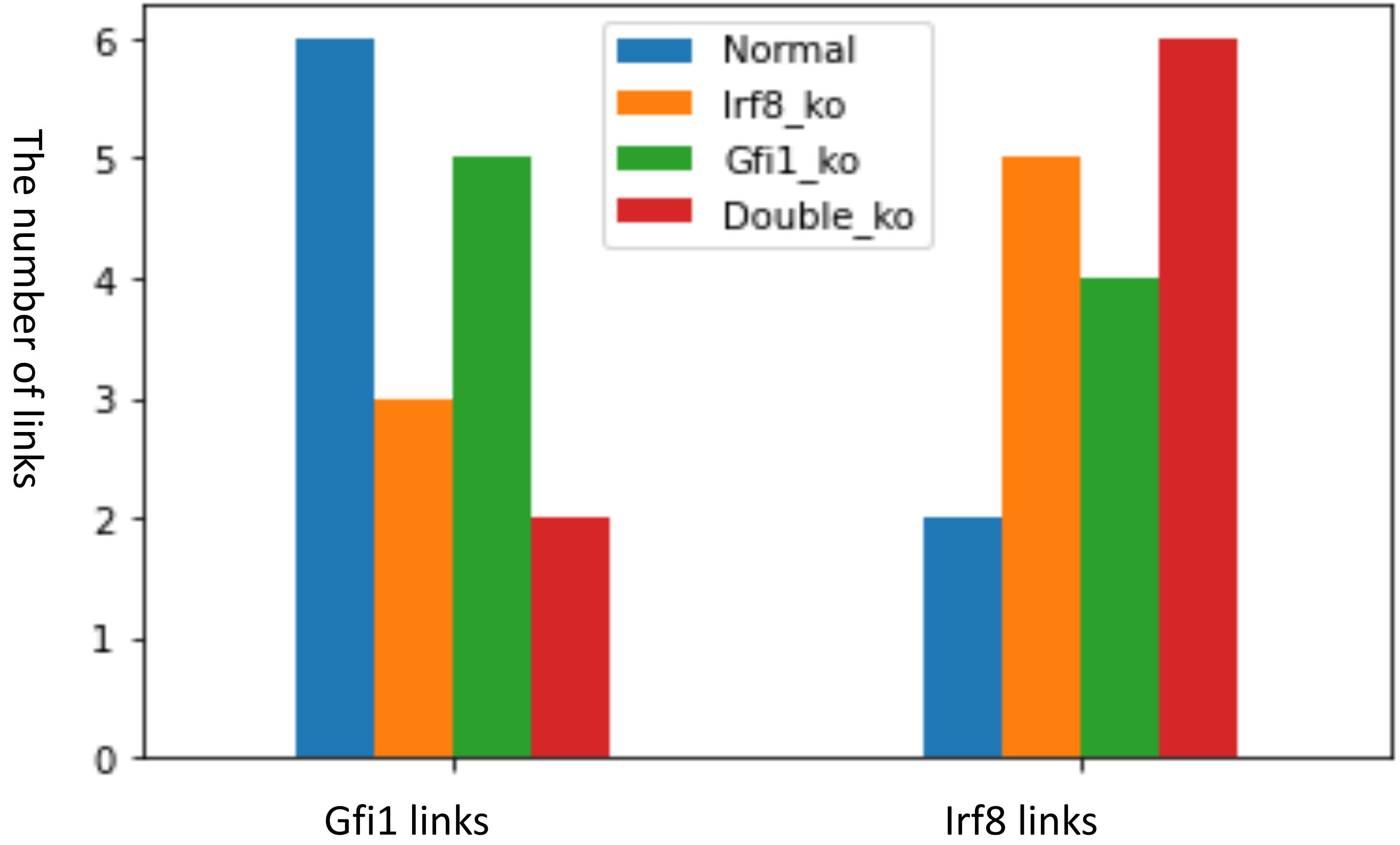
